# Role of gene length in control of human gene expression: chromosome-specific and tissue-specific effects

**DOI:** 10.1101/2020.04.16.044636

**Authors:** Jay C. Brown

## Abstract

**Background:** This study was carried out to pursue the observation that the level of gene expression is affected by gene length in the genomes of higher vertebrates. As transcription is a time-dependent process, it is expected that gene expression will be inversely related to gene length, and this is found to be the case. Here I describe the results of studies performed with the human genome to test whether the gene length/gene expression linkage is affected by two factors, the chromosome where the gene is located and the tissue where it is expressed.

**Experimental design:** Studies were carried out with a database of 2413 human genes that were divided into short, mid-length and long groups. Each of the 24 human chromosomes was then characterized according to the proportion of each gene length group present. A similar analysis was performed with 19 human tissues. The proportion of short, mid-length and long genes was noted for each tissue.

**Results:** Both chromosome and tissue studies revealed new information about the role of gene length in control of gene expression. Chromosome studies led to the identification of two chromosome populations that differ in the level of short gene expression. Tissue studies support the conclusion that short, highly expressed genes are enriched in tissues that produce protein products that are exported from the host cell.

## Introduction

It is now well-established that gene length is associated with the level of gene expression. A high level of expression is found in short genes while expression is weaker in longer ones [1-3]. The same association is observed in a wide variety of eukaryotic organisms [4-6], and there is a good reason to expect the linkage should exist. As time is required to complete transcription of a pre-mRNA molecule, more short molecules are expected to be completed in the same time as fewer longer ones [7-8]. The expected effect of transcription time is consistent with the experimentally observed higher expression of short genes.

One consequence of the association between gene length and gene expression is that length must exert a measure of control over the level of gene expression. If other factors are the same, then longer genes will be expressed at a lower level than shorter ones. If a long gene is to be expressed at a level higher than that determined by its length, then other mechanisms must be at work to adjust the level. Similarly, non-length mechanisms need to be invoked if a short gene, determined by its length to be highly expressed is found to be expressed at a low level. Expression of a gene may therefore be thought of as a background or default level determined by the gene’s length overlaid by other mechanisms to adjust the level according to the requirements for the gene product. Such additional mechanisms may involve well-studied factors such as CpG islands, epigenetic signaling, promoters, transcription factors and others [9-10].

Here I describe the results of studies designed to explore whether expression of a gene as determined by its length may be affected by: (1) the chromosome on which the gene is located; and (2) the tissue where it is expressed. Chromosome studies seek to determine, for example, whether genes with the same length may vary in their expression depending on the chromosome where the gene is located. Studies with tissues compare the level of gene expression as determined experimentally with the level expected due to gene length.

Beginning with a database of human genes, the genes were divided into three groups that differ in length. Genes in each group were then examined for their presence in the 24 human chromosomes and as they are expressed in 19 human tissues. The results are interpreted to clarify the role of chromosome-specific and tissue-specific effects on the level of gene expression in each gene length group.

## Materials and methods

### Gene database

Studies were performed with a database of 2413 human genes that all have tissue selective or tissue specific expression. Broadly expressed (i.e. housekeeping) genes were excluded from the database to make it easier to identify tissue-specific effects of gene length [11]. Genes initially identified for the database were derived from a GRO-seq analysis of genes expressed in IMR90 cells [12]. Nearly all the unexpressed genes in this cell line were found to have either selective or specific gene expression, and these were accepted into the database. The relatively few broadly expressed genes were identified individually and deleted. The database captures a substantial proportion of genes with biased expression (i.e. tissue selective and tissue specifically expressed genes). Estimates of the number of genes with biased tissue expression are in the range of 15% of the total human genes or ∼3000 genes [9, 13], a value consistent with the view that the database (2413 genes) contains a substantial fraction of biased-expression genes.

Five parameters were accumulated for each database gene: (1) the chromosome, (2) the tissue where expression is the highest, (3) whether expression of the gene is tissue-selective or tissue-specific, (4) the level of expression, and (5) whether the gene length is short, mid-length or long. All information was downloaded from the UCSC Genome Browser human genome version hg38 (https://genome.ucsc.edu). Gene expression in a tissue was scored as “specific” if its expression is 10-fold or higher than expression in the tissue with the next highest expression level. Otherwise the gene was scored as having “selective” expression. The lengths of short, mid-length and long genes were <15kb, 15kb-100kb and >100kb, respectively. All gene database information is shown in the Supplementary information (S1 Table) and can be downloaded.

### Data handling

Data were manipulated with Excel and rendered graphically with SigmaPlot v13.0.

## Results

### Gene database

A total of 2413 genes were employed in the study with the number of database genes per chromosome varying between 259 (chr1) and 29 (chr21). All genes are either expressed specifically in a single tissue or they are expressed in only a subset of the tissues reported in the UCSC Genome Browser. Broadly expressed genes were not included in the database. Of the 2413 genes, 562 are expressed only in a single tissue while 1678 are expressed in some but not all tissues (i.e. tissue selective expression). The remainder (173 genes) are too low in expression to be classified.

Using the gene length categories described above, 518 (21.5%) database genes were found to be short, 1259 (52.2%) mid-length and 636 (27.0%) long (Table 1). When the gene composition was examined with individual chromosomes, a wide range was observed in the proportion of short, mid-range and long genes (Table 1). For instance, among the short genes the range observed was 52.7% (chr19) to 3.7% (chr9). With the long genes the range was 49.3% (chr8) to 3.3% (chr19). This observation suggests short genes are located dis-proportionately in some chromosomes and long genes in others.

**Table 1:** Summary of database genes.

| Chr | All Genes | Short Genes | %Short | Mid Genes | %Mid | Long Genes | %Long |
| --- | --- | --- | --- | --- | --- | --- | --- |
| 1 | 259 | 83 | 32.0 | 139 | 53.7 | 37 | 14.3 |
| 2 | 189 | 50 | 26.5 | 84 | 44.4 | 55 | 29.1 |
| 3 | 126 | 18 | 14.3 | 62 | 49.2 | 46 | 36.5 |
| 4 | 147 | 22 | 15.0 | 79 | 53.7 | 46 | 31.3 |
| 5 | 110 | 13 | 11.8 | 58 | 52.7 | 39 | 35.5 |
| 6 | 151 | 24 | 15.9 | 81 | 53.6 | 46 | 30.5 |
| 7 | 137 | 18 | 13.1 | 70 | 51.1 | 49 | 35.8 |
| 8 | 75 | 3 | 4.0 | 35 | 46.7 | 37 | 49.3 |
| 9 | 54 | 2 | 3.7 | 34 | 63.0 | 18 | 33.3 |
| 10 | 87 | 7 | 8.0 | 50 | 57.5 | 30 | 34.5 |
| 11 | 84 | 20 | 23.8 | 50 | 59.5 | 14 | 16.7 |
| 12 | 108 | 18 | 16.7 | 49 | 45.4 | 41 | 38.0 |
| 13 | 54 | 6 | 11.1 | 30 | 55.6 | 18 | 33.3 |
| 14 | 49 | 2 | 4.1 | 27 | 55.1 | 20 | 40.8 |
| 15 | 58 | 7 | 12.1 | 28 | 48.3 | 23 | 39.7 |
| 16 | 93 | 18 | 19.4 | 58 | 62.4 | 17 | 18.3 |
| 17 | 133 | 47 | 35.3 | 71 | 53.4 | 15 | 11.3 |
| 18 | 63 | 8 | 12.7 | 33 | 52.4 | 22 | 34.9 |
| 19 | 150 | 79 | 52.7 | 66 | 44.0 | 5 | 3.3 |
| 20 | 86 | 27 | 31.4 | 45 | 52.3 | 14 | 16.3 |
| 21 | 29 | 7 | 24.1 | 16 | 55.2 | 6 | 20.7 |
| 22 | 41 | 13 | 31.7 | 23 | 56.1 | 5 | 12.2 |
| 23 | 98 | 18 | 18.4 | 52 | 53.1 | 28 | 28.6 |
| 24 | 32 | 8 | 25.0 | 19 | 59.4 | 5 | 15.6 |
| All chr | 2413 | 518 | 21.5 | 1259 | 52.2 | 636 | 27.0 |

Short Genes <15kb; Mid Length Genes 15kb-100kb; Long Genes >100kb

### Inverse relationship between gene length and gene expression

It was expected that gene length would be inversely related to gene expression in the database genes, and that was found to be the case (Fig. 1). Among the three length classes examined, the mean length varied by more than 12-fold. Substantial overlap in expression level was observed among genes in the three length classes, but a clear relationship in mean group values was evident. The observed overlap is consistent with previous analyses [2].

**Fig. 1:**
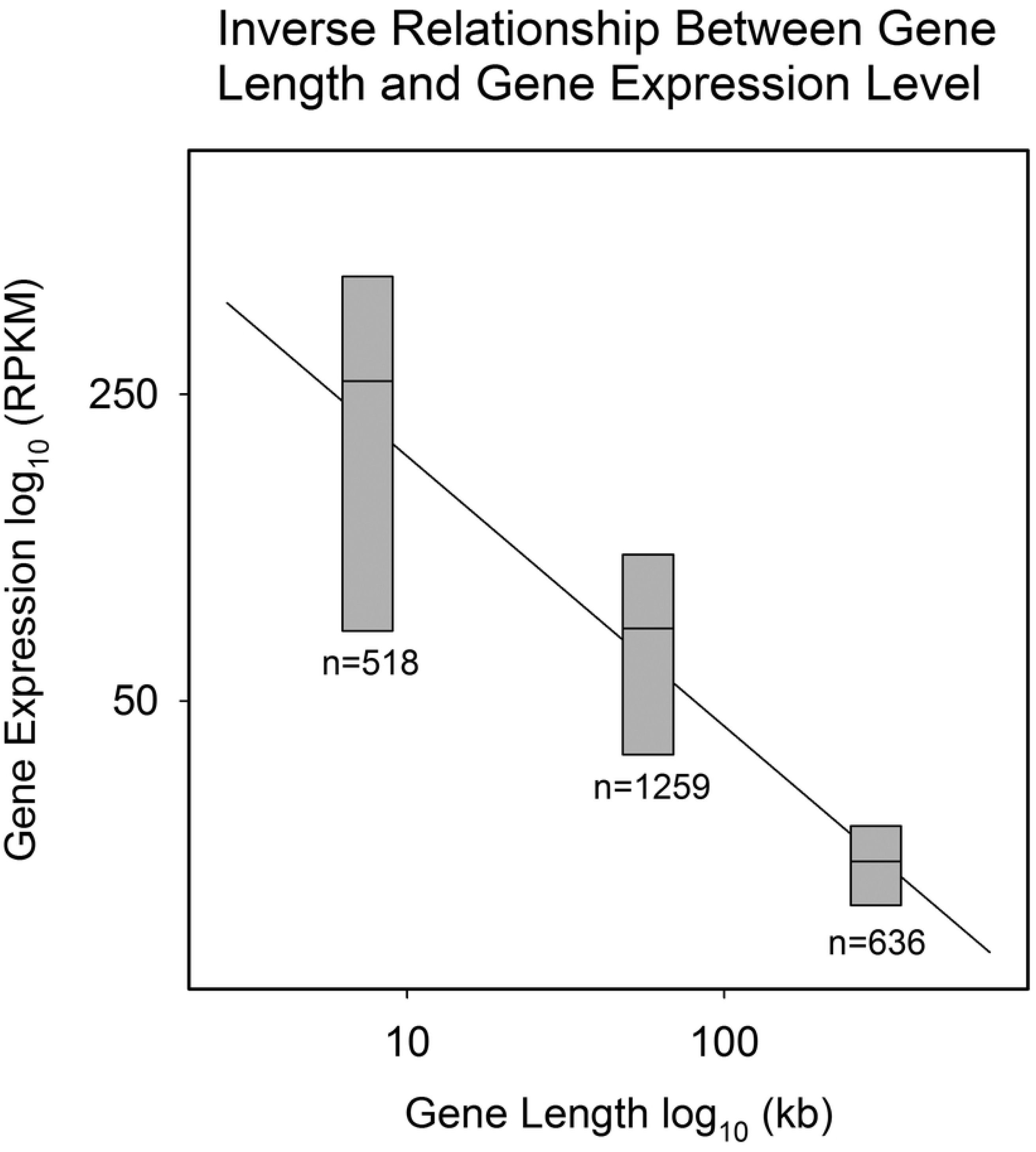
Expression of all database genes in the short, mid-length and long groups. Horizontal lines show the mean expression. Bars indicate the standard error. The length indicated is the mean group length. Note the correlation between shorter genes and higher gene expression.

### Chromosome dependence of gene length

To further characterize the chromosome distribution of long genes, values for the long gene proportion in each chromosome (Table 1) were binned and the count in each bin was plotted against long gene proportion. The results were expected to identify the chromosomes most enriched in long genes. Similar plots were made for mid-length and short genes (Fig. 2).

**Fig. 2:**
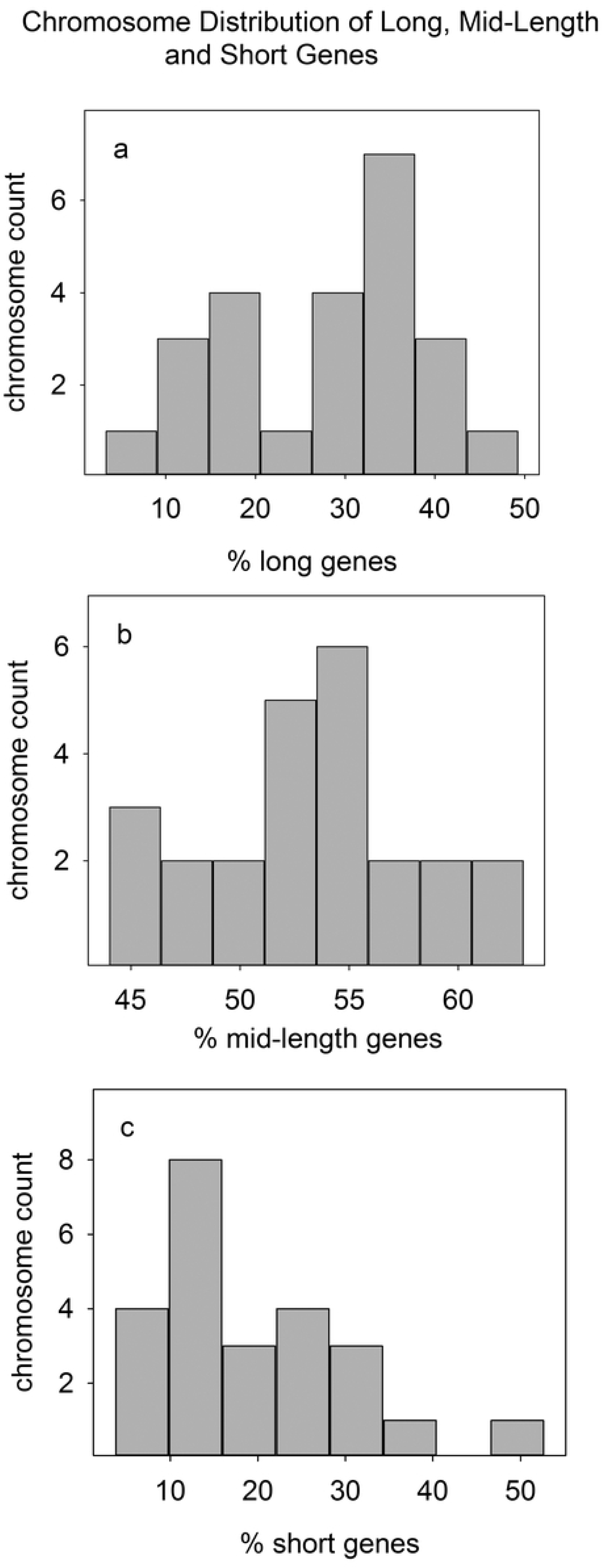
Distribution of short, mid-length and long genes among the 24 human chromosomes. Note that long genes are found in two distinct chromosome groups and many mid-length genes are found in a single, prominent group. The chromosomes in each group are identified in the text.

The results show quite distinct chromosome distributions for the long, mid-length and short gene populations. In the case of the long genes, two populations of chromosomes were observed, one centered at ∼15% long genes and the other at ∼35% (Fig. 2a). Eight chromosomes were found in the low proportion group (1, 11, 16, 17, 19, 20, 22, 24) and fourteen in the high (2-10, 12-15, 18). The existence of distinct chromosome groups indicates that the two differ in their preference for accommodating long genes.

A much narrower distribution was observed in the proportion of mid-length genes. While the proportion of long genes per chromosome extended from ∼2%-49%, in mid-length genes the range was narrower, ∼44%-70% (Fig. 2b). Eleven chromosomes were found in an even tighter distribution from ∼52%-56% mid-length genes (chromosomes 1, 4-6, 13-14, 17-18, 20-21, 23). The result suggests the eleven chromosomes are well adapted to host mid-length genes.

As in the case of long genes, chromosomes were found to have a wide range of values for the proportional count of short genes (range ∼2%-51%; Fig. 2c). The distribution is skewed to lower percentages indicating a preference of some chromosomes for a low proportion of short genes. The distribution is noteworthy for a group of eight chromosomes with ∼12% short genes (Fig. 2c; chromosomes 3-7, 13, 15, 18). Overall, the outcome of chromosomal analysis indicates that human chromosomes differ significantly in the length distribution of the genes they host.

### Chromosomes with high or low proportions of long genes

It was of interest to note the existence of distinct chromosome populations that differ in the proportion of long genes (Fig. 2a). Existence of the two populations suggests that further analysis might lead to identification of factors that affect a chromosome’s gene composition or are affected by it. I examined several possibilities, and the results led to a focus on the expression level of short genes. The mean expression of 198 short genes in the high long gene chromosome population was found to be ∼3-fold higher than the mean of 295 short genes in the low long gene population (Table 2). Control experiments showed no similar increase in expression of long genes in the high compared to the low long gene chromosome populations or in the population of all database genes in the two groups (Table 2).

**Table 2:**
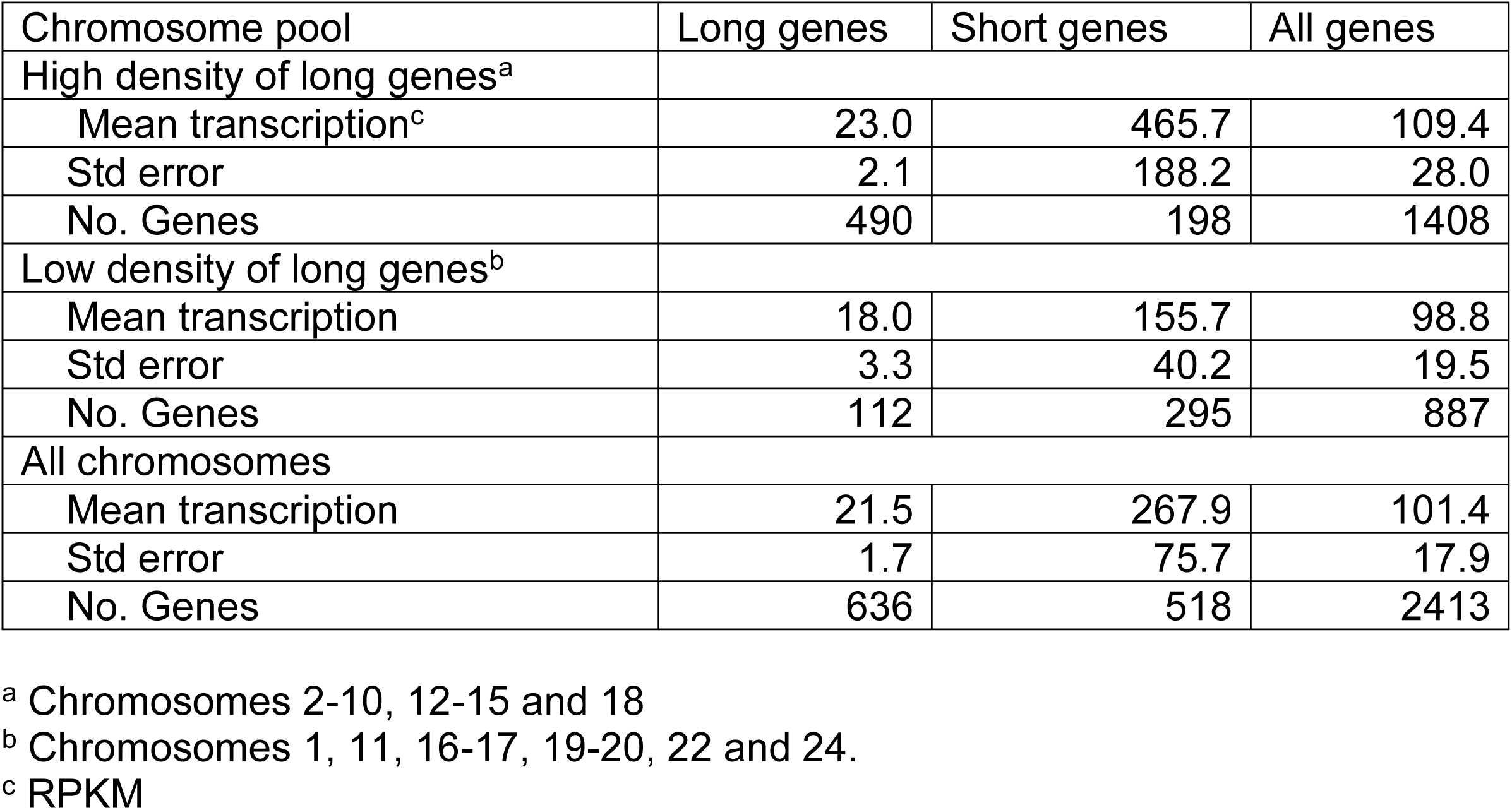
Chromosomes with high compared to low density of long genes.

| Chromosome pool | Long genes | Short genes | All genes |
| --- | --- | --- | --- |
| High density of long genes<sup>a</sup> |  |  |  |
| Mean transcription<sup>c</sup> | 23.0 | 465.7 | 109.4 |
| Std error | 2.1 | 188.2 | 28.0 |
| No. Genes | 490 | 198 | 1408 |
| Low density of long genes<sup>b</sup> |  |  |  |
| Mean transcription | 18.0 | 155.7 | 98.8 |
| Std error | 3.3 | 40.2 | 19.5 |
| No. Genes | 112 | 295 | 887 |
| All chromosomes |  |  |  |
| Mean transcription | 21.5 | 267.9 | 101.4 |
| Std error | 1.7 | 75.7 | 17.9 |
| No. Genes | 636 | 518 | 2413 |

<sup>a</sup> Chromosomes 2-10, 12-15 and 18
<sup>b</sup> Chromosomes 1, 11, 16-17, 19-20, 22 and 24.
<sup>c</sup> RPKM

The above result was not expected and is difficult to interpret. I suggest there may be a chromosome-wide conservation of overall gene expression so that a high number of long, weakly expressed genes is complemented by a population of short, highly expressed ones. Alternatively, chromosomes rich in long genes may be located in regions of the nucleus able to support the high level of transcription characteristic of short genes.

### Tissue dependence of gene length

Beginning with all database genes, each gene was grouped according to its association with one of 19 human tissues, and also with one of the three length groups. The number of genes in each group was then determined and the counts are shown in Table 3.

**Table 3:** Tissue specificity of database genes.

|  | Short Genes |  | Mid Length Genes |  | Long Genes |  | All Genes |  |
| --- | --- | --- | --- | --- | --- | --- | --- | --- |
| Tissue | All<sup>a</sup> | % | All | % | All | % | All | % |
| testis | 141 | 27.2 | 311 | 24.7 | 119 | 18.7 | 571 | 23.7 |
| brain | 67 | 12.9 | 207 | 16.5 | 262 | 41.2 | 536 | 22.2 |
| spleen | 62 | 12.0 | 112 | 8.9 | 21 | 3.3 | 195 | 8.1 |
| liver | 33 | 6.4 | 73 | 5.8 | 11 | 1.7 | 117 | 4.9 |
| skin | 27 | 5.2 | 61 | 4.9 | 9 | 1.4 | 97 | 4.0 |
| sm intestine | 14 | 2.7 | 57 | 4.5 | 9 | 1.4 | 80 | 3.3 |
| esophagus | 14 | 2.7 | 46 | 3.7 | 11 | 1.7 | 71 | 2.9 |
| kidney | 10 | 1.9 | 46 | 3.7 | 10 | 1.6 | 66 | 2.7 |
| muscle | 12 | 2.3 | 33 | 2.6 | 16 | 2.5 | 61 | 2.5 |
| pituitary | 15 | 2.9 | 31 | 2.5 | 16 | 2.5 | 62 | 2.6 |
| adipose | 11 | 2.1 | 26 | 2.1 | 8 | 1.3 | 45 | 1.9 |
| heart | 7 | 1.4 | 25 | 2.0 | 16 | 2.5 | 48 | 2.0 |
| lung | 7 | 1.4 | 25 | 2.0 | 5 | 0.8 | 37 | 1.5 |
| thyroid | 3 | 0.6 | 24 | 1.9 | 29 | 4.6 | 56 | 2.3 |
| adrenal | 4 | 0.8 | 16 | 1.3 | 7 | 1.1 | 27 | 1.1 |
| nerve | 4 | 0.8 | 16 | 1.3 | 15 | 2.4 | 35 | 1.5 |
| pancreas | 14 | 2.7 | 12 | 1.0 | 2 | 0.3 | 28 | 1.2 |
| artery | 8 | 1.5 | 9 | 0.7 | 18 | 2.8 | 35 | 1.5 |
| stomach | 8 | 1.5 | 11 | 0.9 | 2 | 0.3 | 21 | 0.9 |
| other | 57 | 11.0 | 116 | 9.2 | 50 | 7.9 | 223 | 9.2 |
| Total | 518 | 100.0 | 1257 | 100.0 | 636 | 100.0 | 2411 | 100.0 |

<sup>a</sup> All 24 chromosomes.

It was striking to note that the highest number of short and mid-length genes were found in three tissues, testis, brain and spleen (Table 3). Testis and brain were also the top two in number of long genes. I interpret this result to indicate that testis, brain and spleen may require the most genes based on the functions the tissues perform. Other tissues may express fewer genes simply because they don’t need them. The high number of long genes in brain has been noted previously [14].

Tissues were found in four groups based on their distribution of expressed short, mid-length and long genes. In seven of the 19 tissues examined, short genes were the most abundant and long genes the least (Table 3 yellow group). In four tissues, long genes were most abundant (Table 3 green). In the remaining two groups (a) there was little difference among the short, mid-length and long genes or (b) mid-length genes were either the highest or lowest in abundance (Table 3, blue and red groups, respectively). Results for selected tissues are shown graphically in Fig. 3.

**Fig. 3:**
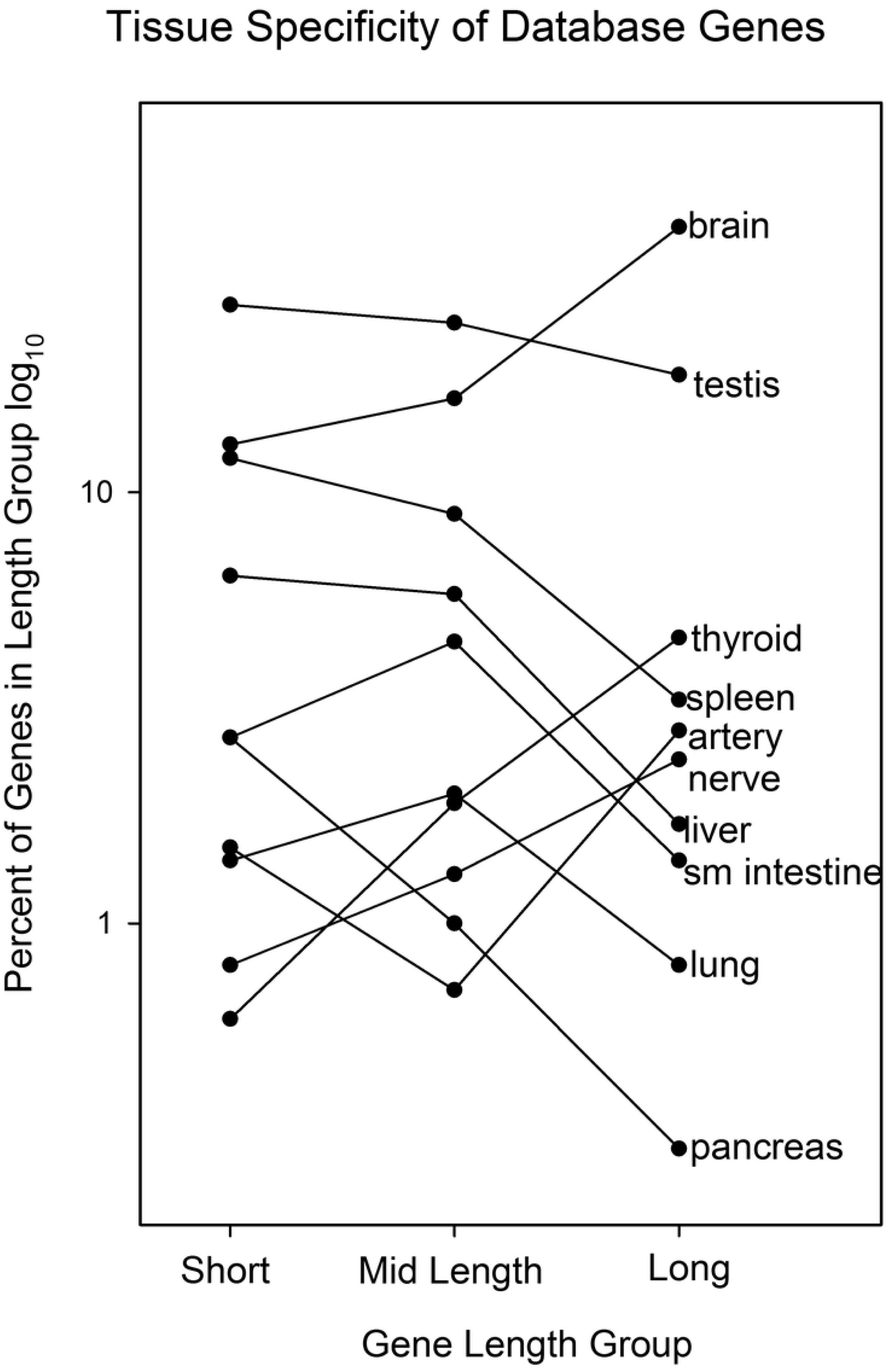
Distribution of database short, mid-length and long genes in ten human tissues. Values shown in plotted form here are selected from the longer list in Table 3. Note that tissues have distinct patters of short, mid-length and long expressed genes.

Reasonable interpretations suggest themselves for some of the results reported in Table 3. For instance, it is expected that tissues involved in synthesizing highly abundant extracellular products would make use of short, highly expressed genes. This is the result observed, for example, with testis, spleen, liver, skin and pancreas. In contrast, brain depends on the function of long proteins involved in processes such as ion uptake, axon guidance and cell adhesion, needs that would be served by expression of long weakly expressed genes (Table 3).

### Tissue-specific effects on transcription level

Tissue specific effects of gene length on transcription level were examined beginning with genes in the same length group. The transcription level of each gene was noted, and the results were compared among the panel of tissues. Controls were provided by the expression level of all database genes in the same length group. Table 4 shows the results obtained with (1) long genes in four different tissues and (2) short genes in brain and liver. The results with long genes show three tissues where the transcription level resembles the control, and one (muscle) where transcription is higher (Table 4). The outcome is said therefore to identify a tissue specific effect of gene length in the case of muscle. The higher expression of muscle genes is suggested to be due to high abundance of muscle tissue where all genes including long ones may need to be expressed at a high level.

**Table 4:** Effect of tissue on expression of genes in the same length group.

| Tissue | Gene Length Group | Mean Transcription (RPKM) | Standard Error | Number Of Genes |
| --- | --- | --- | --- | --- |
| Spleen | Long | 28.2 | 12.7 | 21 |
| Muscle | Long | 82.5 | 29.6 | 16 |
| Pituitary | Long | 16.2 | 5.2 | 16 |
| Thyroid | Long | 39.0 | 17.4 | 29 |
| All tissues | Long | 21.5 | 1.7 | 636 |
| Brain | Short | 47.8 | 11.8 | 66 |
| Liver | Short | 480.5 | 158.2 | 33 |
| All tissues | Short | 267.9 | 75.7 | 518 |

Studies with brain and liver identified tissue specific effects in both cases. Expression of short database genes in liver is higher than the control and lower in brain. The higher expression in liver is suggested to result from the high level of proteins made for export from the liver. Synthesis of abundant exported proteins is expected to require a higher level of gene expression than that needed for use in the home cell only. The opposite situation is observed in brain. As most brain genes encode proteins used in the producing cell, overall gene expression can be low, even in short genes, expected on the basis of their length, to be expressed at a high level.

## Discussion

### Chromosome gene composition

The compositional differences among the human chromosomes shown here supports the view that the chromosomes differ significantly in character (Table 1 and Fig. 2). In chromosome 8, for instance, 49.3% of the genes are in the long group while only 4.0% are in the short. Such distinctions indicate the chromosomes have experienced quite different natural histories before and after they entered the genomes of human progenitor species. Much more now needs to be learned about chromosome evolution before the results in Table 1 and Fig. 2 can be reliably interpreted. For the present, it is safe to assert that the chromosomes are quite individual in their nature and that other aspects of their individuality are likely to emerge in the future [15-16].

Among the most intriguing results of the chromosome composition studies has to do with the identification of two distinct populations of chromosomes, populations that differ in the proportion of long genes encoded (Fig. 2a). Fourteen chromosomes are found in one population and eight in the other. Together the two populations account for 22 of the 24 human chromosomes suggesting the two groups may differ in a binary property that can be in only one state or the other. The two chromosome populations are also of interest because they differ in the expression of short genes (Table 3). Short genes in the high-density long chromosome population are expressed at a higher level than the low. It is tempting to suggest the population of high-density chromosomes just express all genes at a higher level, but this idea is ruled out by the control experiment. High expression is found only among the short and not the long genes (Table 3). The differences between the two chromosome populations may provide the basis for future studies to probe the biochemical properties that underlie their compositional and functional differences.

The chromosome composition studies are also of interest because of the distribution identified among mid-length genes (Fig. 2b). In eleven of the 24 chromosomes the proportion of mid-length genes was found in a narrow distribution between ∼52%-56% of the total genes present. The narrow distribution suggests there is something about the structure or biochemistry of the eleven chromosomes that selects for mid-length gene incorporation and retention in the genome. In contrast, the remaining 13 chromosomes do not demonstrate any similar selection related to mid-length genes (Fig. 2b). Instead, the proportion of mid-length genes is found more evenly distributed over a wider range of mid-length gene content. The result indicates that the selective force that creates a uniform proportion of mid-length genes in eleven chromosomes is lacking in the others creating a wider range of allowed compositional levels.

### Tissue-specific effects

Tissue-specific effects of database genes were examined at two levels, (1) the proportion of short, mid-length and long genes present in a single tissue; and (2) the expression level of genes in the same length class but in different tissues. Studies in the first group address the issue of whereas tissues differ quite significantly in the genes expressed, is this variability accompanied by variability in the distribution of gene lengths? Studies in the second group focus on gene expression. They ask whether genes in the same length group differ in expression when they are present in distinct tissues (Table 4).

Distinctive patterns of expressed gene length were observed in all 19 tissues examined (Table 3 and Fig. 3). Although each tissue pattern was distinctive, four pattern groups could be recognized. They are: (A) most genes are short with decreasing abundance of mid-length and long genes; (B) most genes are long with decreasing abundance of mid-length and short genes; (C) mid-length genes are either highest or lowest in abundance; and (D) short, mid-length and long genes are about equal in abundance. Table 3 shows tissues in the four patterns in yellow, green, red and blue, respectively.

One way to interpret the above pattern groups is to focus on whether a tissue is involved in synthesizing and exporting a protein product. Such export is found in group A tissues including testis, liver and pancreas. These tissues are involved in export of sperm, plasma proteins and digestive enzymes, respectively. As export of such products is expected to require a higher level of gene expression than expression for host cell use only, it is reasonable that short, highly expressed genes should be used. Group B tissues, on the other hand do not synthesize proteins for export. These tissues such as brain and thyroid produce products for local purposes such as neuron function (brain) and small molecule synthesis (thyroid). It is understandable that such tissues should be enriched in long, weakly expressed genes as reported here (Table 3 and Fig. 3).

It was expected that genes of the same length would be found to have quite different expression levels depending on the tissue examined and this was found to be the case (Table 4). It was considered important, however, to establish this issue with genes and tissues from the same database. Expression of long genes in four tissues establish the point. Mean long gene expression in all four tissues differ from each other and from the mean of all long database genes. A similar result was obtained with short genes from liver and brain. Together the results support the view that tissue specific effects influence expression of genes in the same length group.

### Sensing gene length

The results shown here addressing effects of gene length indicate that there need to be cellular mechanisms able to sense gene length. It would be impossible, for instance, for chromosome 19 to have a high proportion of short genes (52.7%; Table 1) if there were no mechanism for the cell to sense short genes or something that correlates with gene shortness. It is relevant therefore that mechanisms of the expected type have been reported in the case of long genes. Studies with mouse, for instance, have demonstrated that the Mecp2 gene selectively downregulates long gene expression [17]. Mecp2 is found to act by binding selectively to methylated CA-containing DNA sites in long genes. Depletion of the same activity in the human homolog, MECP2, is found to cause Rett syndrome [18].

Two other instances have been reported [19-20]. The protein encoded by the human SFPQ gene is found to bind to introns in long genes ensuring that transcription will proceed to the end of the gene. Similarly, human topoisomerases TOPI and TOPII have been shown to facilitate the transcription of long genes in neurons [20]. In view of the results reported here it is reasonable to suggest that there may be similar cellular systems to recognize short and mid-length genes.

## Acknowledgements

I gratefully acknowledge Forde Upshur, Ava Roth and Karsten Siller for advice on computational aspects of this project.

## References

1. Castillo-Davis CI, Mekhedov SL, Hartl DL, Koonin EV, Kondrashov FA. Selection for short introns in highly expressed genes. Nat Genet. 2002;31(4):415–8.

2. Urrutia AO, Hurst LD. The signature of selection mediated by expression on human genes. Genome Res. 2003;13(10):2260–4.

3. Chiaromonte F, Miller W, Bouhassira EE. Gene length and proximity to neighbors affect genome-wide expression levels. Genome Res. 2003;13(12):2602–8.

4. Munoz ET, Bogarad LD, Deem MW. Microarray and EST database estimates of mRNA expression levels differ: the protein length versus expression curve for C. elegans. BMC Genomics. 2004;5(1):30.

5. Grishkevich V, Yanai I. Gene length and expression level shape genomic novelties. Genome Res. 2014;24(9):1497–503.

6. Nie H, Crooijmans RP, Lammers A, van Schothorst EM, Keijer J, Neerincx PB, et al. Gene expression in chicken reveals correlation with structural genomic features and conserved patterns of transcription in the terrestrial vertebrates. PLoS One. 2010;5(8):e11990.

7. Veloso A, Kirkconnell KS, Magnuson B, Biewen B, Paulsen MT, Wilson TE, et al. Rate of elongation by RNA polymerase II is associated with specific gene features and epigenetic modifications. Genome Res. 2014;24(6):896–905.

8. Danko CG, Hah N, Luo X, Martins AL, Core L, Lis JT, et al. Signaling pathways differentially affect RNA polymerase II initiation, pausing, and elongation rate in cells. Mol Cell. 2013;50(2):212–22.

9. Deaton AM, Bird A. CpG islands and the regulation of transcription. Genes Dev. 2011;25(10):1010–22.

10. Portela A, Esteller M. Epigenetic modifications and human disease. Nat Biotechnol. 2010;28(10):1057–68.

11. Zhu J, He F, Hu S, Yu J. On the nature of human housekeeping genes. Trends Genet. 2008;24(10):481–4.

12. Core LJ, Waterfall JJ, Lis JT. Nascent RNA sequencing reveals widespread pausing and divergent initiation at human promoters. Science. 2008;322(5909):1845–8.

13. Brown JC. Control of human testis-specific gene expression. PLoS One. 2019;14(9):e0215184.

14. Barbash S, Sakmar TP. Length-dependent gene misexpression is associated with Alzheimer’s disease progression. Sci Rep. 2017;7(1):190.

15. Capozzi O, Purgato S, D’Addabbo P, Archidiacono N, Battaglia P, Baroncini A, et al. Evolutionary descent of a human chromosome 6 neocentromere: a jump back to 17 million years ago. Genome Res. 2009;19(5):778–84.

16. Murphy WJ, Fronicke L, O’Brien SJ, Stanyon R. The origin of human chromosome 1 and its homologs in placental mammals. Genome Res. 2003;13(8):1880–8.

17. Gabel HW, Kinde B, Stroud H, Gilbert CS, Harmin DA, Kastan NR, et al. Disruption of DNA-methylation-dependent long gene repression in Rett syndrome. Nature. 2015;522(7554):89–93.

18. Chahrour M, Zoghbi HY. The story of Rett syndrome: from clinic to neurobiology. Neuron. 2007;56(3):422–37.

19. Takeuchi A, Iida K, Tsubota T, Hosokawa M, Denawa M, Brown JB, et al. Loss of Sfpq causes long-gene transcriptopathy in the brain. Cell Rep. 2018;23(5):1326–41.

20. King IF, Yandava CN, Mabb AM, Hsiao JS, Huang HS, Pearson BL, et al. Topoisomerases facilitate transcription of long genes linked to autism. Nature. 2013;501(7465):58–62.

